# A Protease-Cleavable iNOS-Inhibitor Polymeric Prodrug Designed for Controlled Modulation of Nitric Oxide

**DOI:** 10.64898/2026.06.23.733308

**Authors:** Houman Alimoradi, Azam Panahpour, Anita Fallah, Christine Delporte

**Affiliations:** Laboratory of Pathophysiological and Nutritional Biochemistry (LPNB), Faculty of Medicine, Université Libre de Bruxelles, Brussels, Belgium; Department of Molecular Medicine, Faculty of Advanced Medical Sciences, Tabriz University of Medical Sciences, Tabriz, Iran

**Keywords:** Nitric oxide, iNOS, Prodrug, Protease-responsive

## Abstract

Inducible nitric oxide synthase (iNOS) is frequently overexpressed in inflammatory disorders and solid tumors, where sustained nitric oxide (NO) production promotes angiogenesis, tumor progression, and resistance to therapy. Despite promising preclinical results, the clinical translation of iNOS inhibitors remains limited by poor tumor selectivity, rapid systemic clearance, and off-target toxicities. To address these challenges, we developed a protease-responsive polymeric iNOS-inhibiting prodrug (ProCIP) designed for localized activation within protease-rich pathological microenvironments. ProCIP was synthesized from poly(ethylene glycol)-poly(L-glutamate) and functionalized with amidine-based iNOS inhibitory moieties. The resulting cationic polymer readily formed nanoscale polyionic complexes with anionic polymers or molecules. In cell-free assays, enzymatic activation of ProCIP resulted in a significant reduction in iNOS activity, whereas non-activated nanoparticles showed minimal inhibition. Cellular studies confirmed efficient nanoparticle uptake by RAW264.7 macrophages and revealed a significant reduction in intracellular NO levels in lipopolysaccharide-stimulated cells. These findings demonstrate that ProCIP enables protease-triggered iNOS inhibition and localized NO regulation, offering a promising strategy for improving the safety and efficacy of iNOS-targeted therapies in cancer and other inflammatory diseases.

## 1. Introduction

Nitric oxide (NO) is a versatile biological mediator that regulates numerous physiological functions, and disturbances in its production are implicated in a wide range of diseases [1]. The effects of NO depend not only on its concentration but also on the duration and location of its generation. In cancer, inducible nitric oxide synthase (iNOS) is a major source of sustained NO production and exerts diverse biological effects that vary according to tumor type and microenvironmental context [1]. Overexpression of iNOS has been reported in multiple aggressive cancers, leading to prolong NO exposure within the tumor microenvironment (TME) [2, 3]. Although excessive NO concentrations can trigger cytotoxic responses, including DNA damage and apoptosis, the lower yet persistent levels commonly observed in tumors are frequently associated with enhanced proliferation, angiogenesis, immune evasion, and therapeutic resistance [1].

In colorectal cancer (CRC), increased iNOS expression has been linked to disease progression and adverse patient outcomes [3, 4]. Several clinical investigations have demonstrated a positive association between iNOS expression and metastatic spread to regional lymph nodes, as well as reduced survival rates [5, 6]. Importantly, the prognostic value of iNOS appears to be independent of conventional clinicopathological indicators. Patients with iNOS-positive tumors exhibit a substantially greater risk of disease-specific mortality than those lacking detectable iNOS expression, highlighting the contribution of NO-driven signaling to CRC progression [5].

The therapeutic relevance of iNOS has prompted efforts to evaluate its inhibition as an adjuvant anticancer strategy. Experimental and clinical studies indicate that suppression of iNOS activity can improve the response to taxane-based treatments, including paclitaxel (PTX), through modulation of pathways involved in tumor survival and dissemination [7]. In preclinical CRC models, iNOS inhibitors have also demonstrated synergistic effects when combined with standard chemotherapeutic regimens, resulting in reduced tumor growth and attenuation of pro-tumorigenic signaling within the TME [8].

Despite these promising findings, the clinical application of iNOS inhibitors remains limited. A major challenge arises from the multifaceted roles of NO in normal physiology and disease, making selective modulation difficult. Furthermore, currently available inhibitors often lack complete isoform specificity, increasing the risk of adverse effects such as hypertension caused by unintended inhibition of endothelial nitric oxide synthase (eNOS) [7, 9]. Another significant limitation is the unfavorable pharmacokinetic profile of potent inhibitors such as 1400W, which exhibit rapid systemic elimination and poor accumulation at tumor sites, thereby restricting their therapeutic effectiveness [10-12].

To overcome these barriers, we engineered a polymeric prodrug platform capable of delivering an iNOS inhibitor selectively within the colorectal tumor microenvironment. The system was designed to respond to elevated protease activity, a hallmark of many colorectal tumors, enabling site-specific activation and controlled release of the inhibitor. By restricting drug liberation to the tumor milieu, this approach aims to enhance therapeutic efficacy while minimizing systemic toxicity and off-target pharmacological effects.

### 2. Materials and Methods

Unless otherwise specified, all chemicals were used without further purification. Reagents were obtained from commercial suppliers, predominantly TCI Europe (Zwijndrecht, Belgium) and Sigma-Aldrich (St. Louis, MO, USA). Solvents employed for synthesis and sample preparation were purchased from Thermo Fisher Scientific (Waltham, MA, USA) or Merck (Carlsbad, CA, USA). Structural characterization of synthesized compounds was performed by proton nuclear magnetic resonance (^1^H NMR) spectroscopy using a 400 MHz JEOL instrument (Tokyo, Japan). Spectral acquisition and analysis were carried out with MestReNova software (version 15, CIREM license). Functional group analysis was conducted by Fourier-transform infrared spectroscopy (FTIR) on a Bruker Alpha II spectrometer (Bruker Optics GmbH & Co. KG, Ettlingen, Germany) operating in attenuated total reflectance (ATR) mode. Spectra were acquired between 4000 and 500 cm□^1^ at a resolution of 1 cm□^1^, with 32 scans collected for each sample.

Nanoparticle (NP) surface charge was determined by zeta potential measurements using a Zetasizer Ultra instrument (Malvern Instruments Ltd., Malvern, UK). Optical characterization was performed by UV-visible spectroscopy on a PerkinElmer LAMBDA™ 25/35 spectrophotometer (PerkinElmer, Springfield, IL, USA).

### 2.1. Synthesis of ProCIP

#### 2.1.1. Synthesis of PEG-Poly(L-glutamate) (PEG-PGA)

PEG-poly(L-glutamate) (PEG-PGA) was synthesized via ring-opening polymerization (ROP) of γ-benzyl-L-glutamate N-carboxyanhydride (BLG-NCA), followed by catalytic hydrogenolysis of the benzyl protecting groups (Scheme 1). Briefly, methoxy poly(ethylene glycol) amine (MeO-PEG-NH□, 5 kDa) was used as a macroinitiator for the polymerization. BLG-NCA (2.1 g, 8 mmol) was dissolved in anhydrous dimethylformamide (DMF, 10 mL), and a solution of MeO-PEG-NH□ (2.4 g, 0.4 mmol) in anhydrous DMF (10 mL) was added under a nitrogen atmosphere, corresponding to a monomer-to-initiator feed ratio of 20:1. The reaction mixture was stirred at 40 °C under anhydrous and inert conditions for 48 h. Upon completion of the polymerization, the reaction mixture was precipitated into excess cold diethyl ether. The resulting polymer was redissolved in a minimal volume of DMF (1-2 mL) and reprecipitated into cold diethyl ether twice more to remove residual monomer and solvent. The purified polymer was collected by filtration and dried under vacuum at 40 °C to afford Compound **1** as a white powder.

To obtain PEG-poly(L-glutamic acid) (PEG-PGA, Compound **2**), the benzyl protecting groups were removed by catalytic hydrogenolysis. **1** was dissolved in dry tetrahydrofuran (THF) and treated with 10 wt% palladium on carbon (Pd/C) under a hydrogen atmosphere at room temperature for 24 h. Upon completion of the reaction, the catalyst was removed by centrifugation followed by filtration through a membrane filter. The filtrate was concentrated under reduced pressure and precipitated into methanol. The resulting polymer was collected and dried under vacuum to afford PEG-PGA as a white solid.

**Scheme 1.**
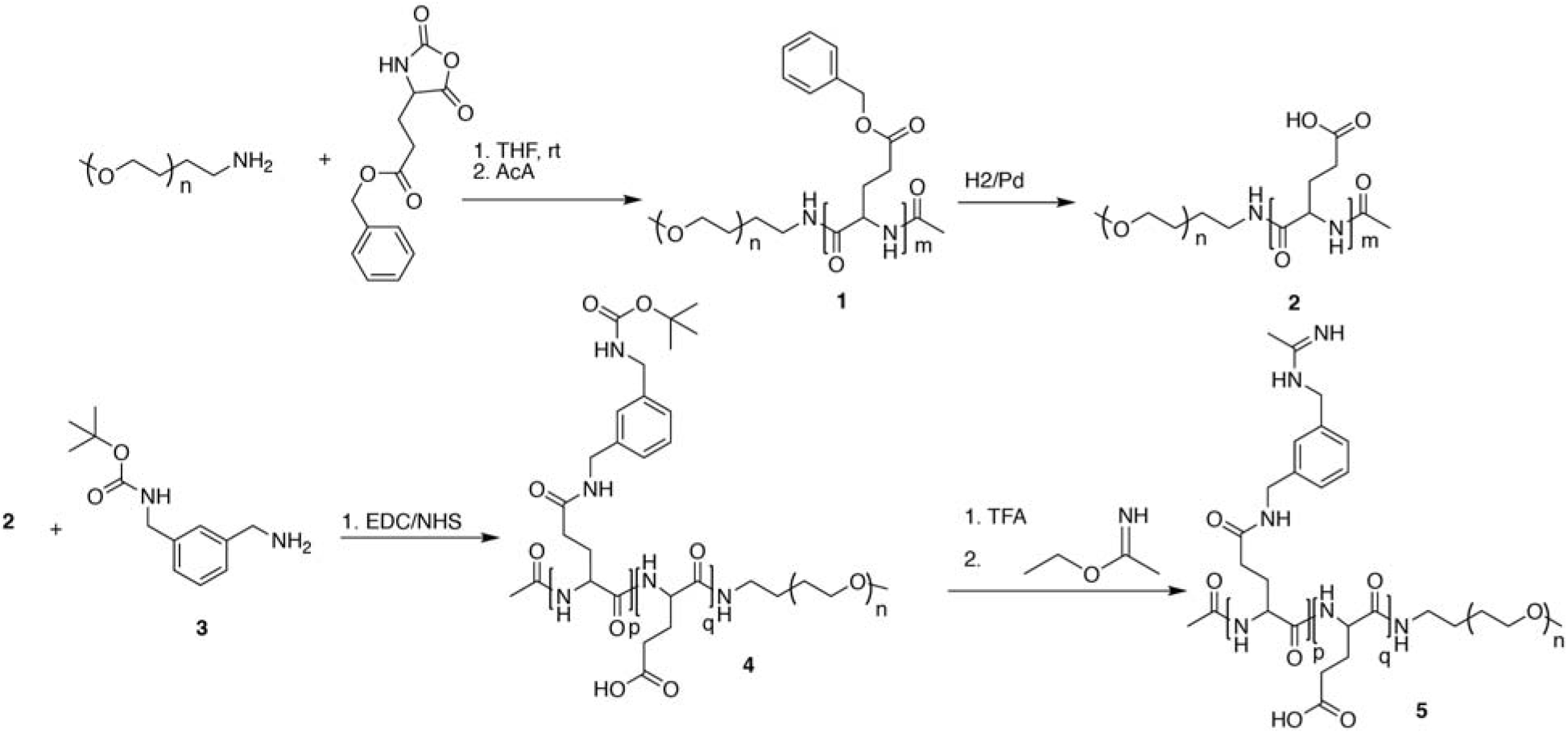
Mechanism of synthesis of ProCIP (**5**).

#### 2.1.2. Conjugation of aminomethyl benzene

Compound **4** was synthesized by EDC/NHS-mediated coupling of PEG-poly(L-glutamic acid) (PEG-PGA) with 1-(N-Boc-aminomethyl)-3-(aminomethyl)benzene (Compound **3**). Briefly, PEG-PGA (2.0 g) was dissolved in anhydrous DMF (10 mL), followed by the addition of EDC·HCl (0.46 g) and NHS (0.62 g). The mixture was stirred at room temperature for 15 min to activate the pendant carboxyl groups of the polymer. In a separate vial, compound **3** (1.0 g, 4.2 mmol) was dissolved in DMF (2 mL) and added dropwise to the activated polymer solution. The reaction mixture was stirred at room temperature for 24 h under a nitrogen atmosphere.

The crude reaction mixture was then added dropwise to cold diethyl ether (100 mL) to precipitate the polymer conjugate. The precipitate was collected by centrifugation (1000 × g), redissolved in a minimal volume of DMF (1-2 mL), and reprecipitated into cold diethyl ether. This dissolution-precipitation cycle was repeated two additional times to remove unreacted reagents and low-molecular-weight byproducts. The purified product was isolated by centrifugation and dried under vacuum to afford Compound **4** as a white solid.

#### 2.1.3. Boc deprotection and formation of imidate-modified polymer

To remove the tert-butyloxycarbonyl (Boc) protecting groups, **4** (2.0 g) was dissolved in dichloromethane (DCM), and trifluoroacetic acid (TFA) was added dropwise to afford a 30% (v/v) TFA/DCM solution. The reaction mixture was stirred at room temperature for 4 h. Upon completion of the deprotection, the solvents were removed under a stream of nitrogen. The residue was redissolved in methanol and concentrated under reduced pressure to facilitate removal of residual TFA, yielding the corresponding amine-functionalized intermediate.

The deprotected polymer was subsequently dissolved in ethanol (15 mL), and triethylamine was added to adjust the solution to basic conditions. Ethyl acetimidate hydrochloride (1.23 g, 10 mmol), dissolved in ethanol (5 mL), was then added dropwise to the reaction mixture. The resulting solution was stirred at room temperature for 24 h to afford the amidine-functionalized polymer.

Following completion of the reaction, the mixture was transferred to dialysis tubing and dialyzed against deionized water for 48 h with regular water replacement. The purified product was recovered by lyophilization to yield Compound **5** (ProCIP) as a white solid.

### 2.2. iNOS inhibition assay

iNOS activity was determined by measuring NO generation using a Griess-based colorimetric assay. The enzymatic reaction was performed in 50 mM HEPES buffer containing iNOS (3.6 U mL□^1^, Sigma-Aldrich), NADPH (300 μM), magnesium acetate (1.0 mM), BH4 (180 μM), and DTT (180 μM) [13]. Reaction mixtures were incubated at 37 °C for 45 min to allow NO production. Following incubation, aliquots were treated with Griess reagents according to the manufacturer’s protocol, resulting in the formation of a chromophoric azo compound proportional to the amount of NO-derived nitrite present in the sample. Absorbance was recorded at 540 nm using a spectrophotometer. Nitrite concentrations were determined from a standard calibration curve generated with sodium nitrite, and iNOS activity was expressed as relative NO production.

### 2.4. Cell culture and treatment

Cell culture experiments were carried out using media and biological reagents supplied by Thermo Fisher Scientific. RAW264.7 macrophages (ATCC, Manassas, VA, USA) were maintained in DMEM/F12 supplemented with 5% heat-inactivated FBS, 100 U mL□^1^ penicillin-streptomycin, and 4 mM L-glutamine. Cells were grown in a humidified atmosphere containing 5% CO□ at 37 °C and subcultured according to standard procedures. The inhibitory effects of the NPs (made of CMC and ProCIP), was assessed in the cells against LPS-induced NO production. LPS treatment was carried out at the concentration of 0.5 μg/mL for 24 h, to induce iNOS expression and NO synthesis in the macrophages.

### 2.5. Evaluation of NO production

The ability of the test compounds to modulate intracellular NO levels was investigated following 24 h exposure periods. After treatment, cells were incubated with DAF-2DA (4 μM) prepared in serum-free DMEM for 30 min at 37 °C to enable detection of intracellular NO. Cells were subsequently counterstained with DAPI for 10 min to visualize cell nuclei. Following staining, excess dye was removed by washing the cells twice with PBS. Fluorescence images were acquired using a LSM900 confocal microscope equipped with Airyscan (Carl Zeiss, Oberkochen, Germany) under identical acquisition settings for all experimental groups [14-16]. For flow cytometric quantification of intracellular NO, cells were detached, collected as single-cell suspensions, and stained with DAF-2DA using the same concentration and incubation conditions described above. Fluorescence intensity was then measured using a FACScan flow cytometer (Becton Dickinson, NJ, USA) and analyzed using FlowJo software (Ashland, OR, USA), and the resulting signal was used as an indicator of intracellular NO production.

### 2.6. Statistical analysis

Data are reported as mean ± SD. Statistical comparisons were conducted using GraphPad Prism (version 9.0; GraphPad Software). Group differences were analyzed by one-way ANOVA followed by Tukey’s post hoc test. Values of p < 0.05 were considered statistically significant.

## 3. Results and Discussion

### 3.1. Synthes is of polymeric prodrug

Pro CIP was synthesized following the synthetic pathway illustrated in Scheme 1. Initially, PEG-PPG was prepared using a well-established ring-opening polymerization (ROP) method, as previously described [17]. The polymerization of BLG-NCA was initiated by a 5Kda PEGAmine which was designed to prevent biological interactions with the resulting prodrug. HNMR of PEG-PPG shows the presence of the resonance of aromatic protons at δ ∼ 7.3 ppm, benzylic protons at 5.0 ppm, α-proton of gluthamates at δ ∼ 4.01 ppm and the broad peaks between 1.9-2.4 ppm which belong to -CH2- of gluthamate side changes. Benzylic groups were deprotected using H□ gas on Pd/C, and this was confirmed by the disappearance of aromatic protons and benzylic -CH□-(Figure 1a). PEG-PGA was then coupled with tert-butyl 3-(aminomethyl)benzylcarbamate using EDC/NHS. The appearance of significant tert-butyl peaks of BOC (δ ∼1.4 ppm), aromatic protons, and benzylic -CH□-in the aminomethyl (-CH□NH□) and N-Boc-aminomethyl (-CH□NHBoc) substituents around 4.2 ppm confirmed the coupling reaction. To prepare ProCIP, Boc groups were initially deprotected with trifluoroacetic acid (TFA), followed by reaction with ethyl acetimidate in ethanol [18]. This resulted in the disappearance of tert-butyl protons and the appearance of an acetimidate peak at 2.2 ppm (Figure 1a). FT-IR spectroscopy further confirmed the successful synthesis of ProCIP (Figure 1b). A broad absorption band extending from approximately 3600 to 3100 cm□^1^ was observed and is attributed to overlapping O-H stretching vibrations of carboxylic acid groups and N-H stretching vibrations of amide and acetimidate functionalities. The characteristic aliphatic C-H stretching vibrations appeared around 2950-2850 cm□^1^. A strong absorption band at approximately 1660 cm□^1^ corresponds to the amide I carbonyl (C=O) stretching vibration of the polypeptide backbone and pendant carboxylic acid groups. The band observed near 1540-1560 cm□^1^ can be assigned to acetimidate (C=N stretching) and aromatic ring vibrations were detected in the region of 1450-1500 cm□^1^. The intense band centered around 1100-1150 cm□^1^ is characteristic of ether (C-O-C) stretching vibrations of the PEG segment, confirming the presence of the PEG backbone. Together, these characteristic absorptions support the successful synthesis of ProCIP.

**Figure 1.**
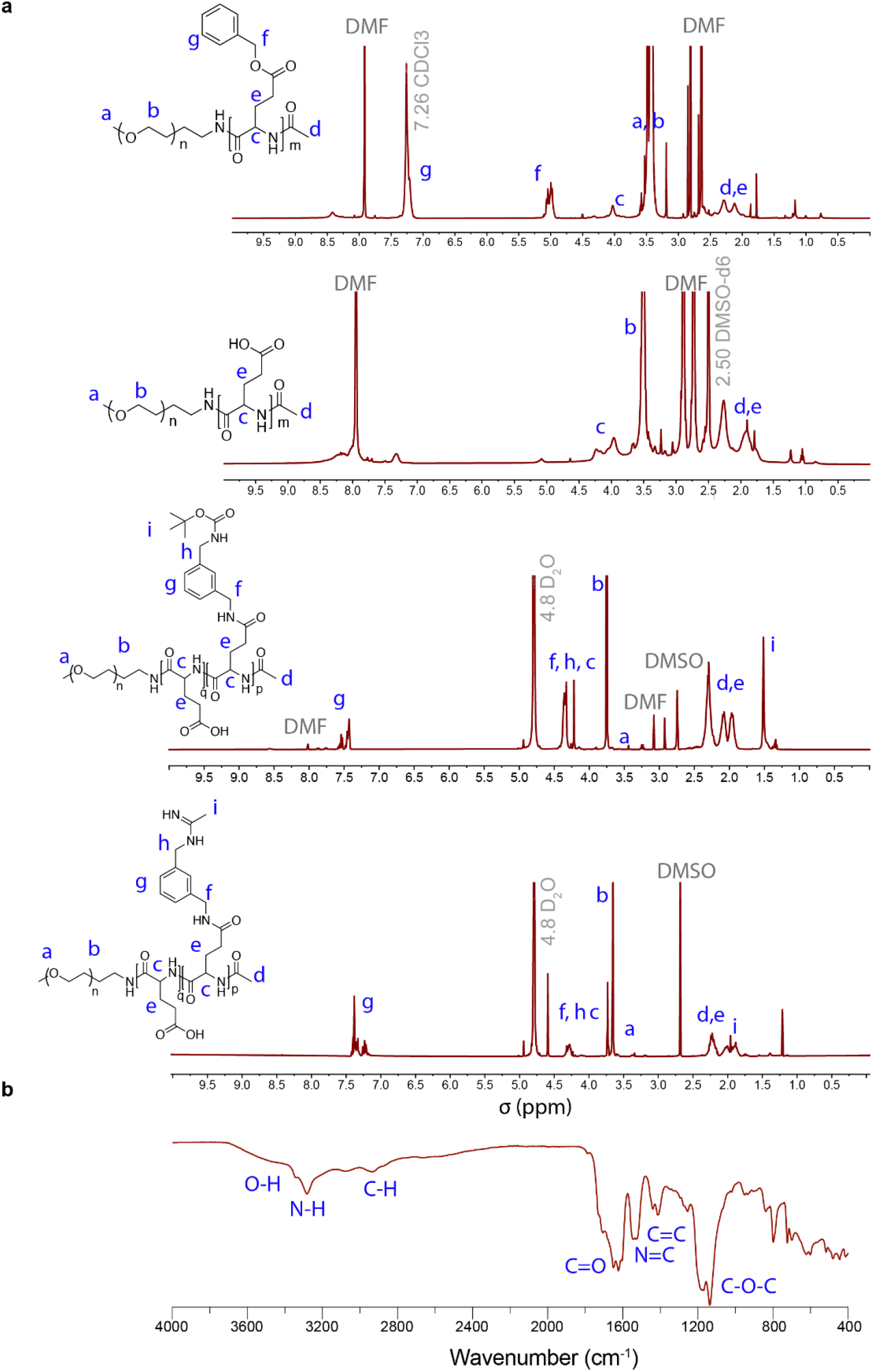
Characterization of ProCIP synthesis using ^1^HNMR (**a**) and FT-IR (**b**).

### 3.2. Formulation with anionic molecules

Polyionic complexes have attracted considerable attention as versatile drug delivery systems due to their straightforward, solvent-free preparation, excellent biocompatibility, and highly tunable physicochemical properties [19]. The electrostatic interactions between oppositely charged polymers or molecules facilitate the efficient encapsulation of a broad range of therapeutic agents. Furthermore, drug release can be finely controlled by environmental stimuli, including pH, ionic strength, and enzymatic activity [19]. Such responsiveness enables the design of delivery platforms capable of adapting to specific physiological microenvironments, thereby improving site-selective drug release while minimizing off-target effects.

ProCIP exhibits a positively charged surface, with an average zeta potential of +6.0 ± 1.2 mV. Upon interaction with anionic polymers, such as carboxymethyl cellulose (CMC), or negatively charged hydrophobic molecules containing acidic functional groups (e.g., Eosin Y and Cy5.5), ProCIP readily forms nanoscale polyelectrolyte complexes. The addition of CMC to a vigorously stirred ProCIP suspension induced charge neutralization and particle assembly, resulting in a marked increase in particle size (Figure 2). The resulting NPs displayed an average hydrodynamic diameter of 179 ± 85 nm and a near-neutral surface charge, with a zeta potential of −0.5 ± 3.6 mV (Figure 2).

**Figure 2.**
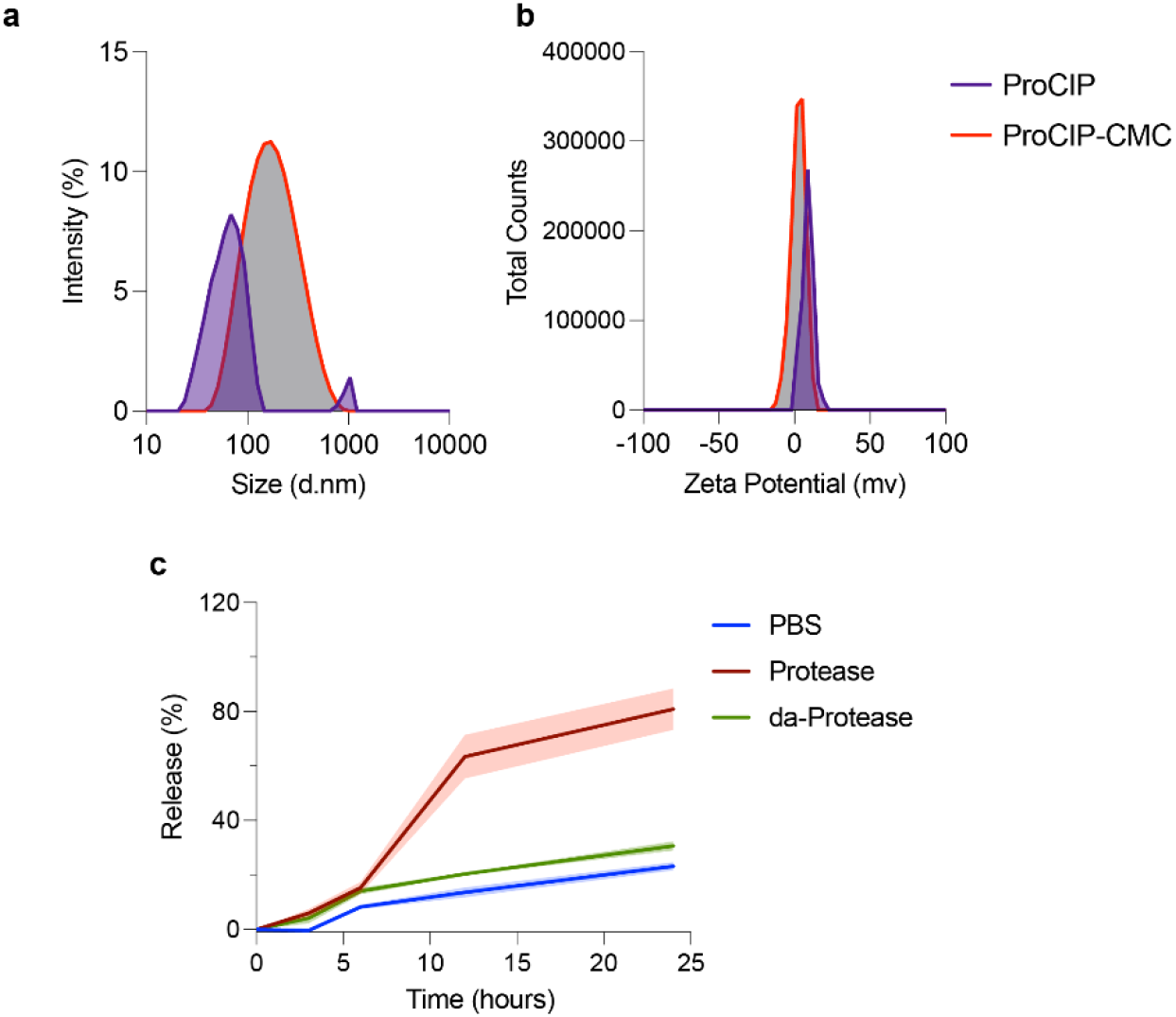
Characterization of ProCIP synthesis using ^1^HNMR. **a**. Representative histogram of NP size distribution measured by DLS using a 1 mg/mL NP solution in distilled water. **b**. Zeta potential of ProCIP and NPs made of ProCIP-CMC measured in 10 mM NaCl solution. **c**. Release of eosin from the NPs made ProCIP and eosin. Data are expressed as mean ± SD (n=3).

The NPs exhibited a sustained release profile under physiological conditions. Approximately 13.5% of the encapsulated payload was released during the first 12 h, increasing to 32.9% after 72 h. Supplementation with 1 mg mL□^1^ bovine pancreatic protease resulted in a 4.7-fold enhancement of payload release within the initial 12 h. Because enzymes, like many proteins, can exert surfactant-like effects that may influence NP stability [20], heat-inactivated proteases at the same concentration were employed as a control. Under these conditions, eosin release increased by 6.9% after 12 h, indicating that the substantially enhanced release observed in the presence of active proteases was primarily driven by enzymatic degradation of the NP matrix rather than nonspecific protein-mediated destabilization.

### 3.3. The effects of NPs on iNOS inhibition

NPs made of ProCIP and CMC did not show any effect on the enzymatic activity of iNOS at 50□µg/mL. However, when the NPs were exposed to bovine pancreas proteases for 24h followed by heat deactivation of the proteases, a 76% reduction in iNOS activity was observed, indicating a hydrolysis-dependent activation (Figure 3A).

**Figure 3.**
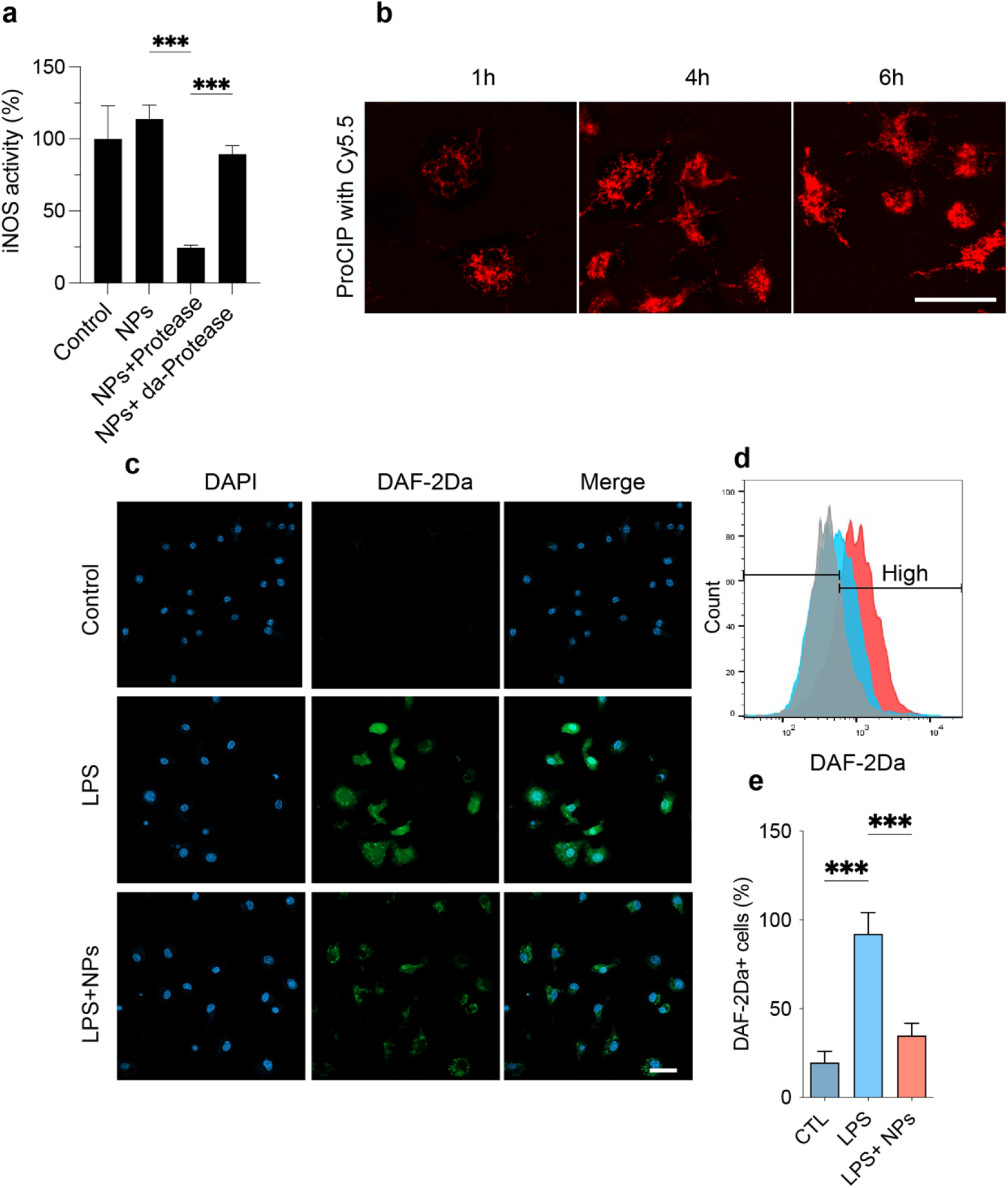
Effects of ProCIP on iNOS activity. **a**. iNOS enzymatic inhibition assay of 50 μg/mL NPs made of ProCIP and CMC. **b**. Endocytosis of NPs made of ProCIP and a fluorescent dye (Cy5.5). **c**. Confocal microscopy images of RAW264.7cells stained with DAF-2Da and incubated with LPS (0.5 μg/mL), and then without or with 30 μg/mL NPs made of ProCIP-CMC. **d**. Representative flow cytometry histogram of NO levels in DAF-2Da-stained RAW264.7 cells following incubation with the treatments. **e**. Quantification of high-fluorescence DAF-2Da-positive RAW264.7. The comparison was done by one-way ANOVA with Tukey’s post hoc. Data are expressed as mean ± SD (n=3). ***p < 0.001 vs. control. Scale bar is 20μm.

To evaluate the activity of the NPs in RAW264.7 macrophages displaying high iNOS expression, we first assessed cellular uptake by formulating NPs composed of ProCIP and the fluorescent dye Cy5.5. The NPs were internalized by cells starting from 1h incubation, and their intracellular concentration increased over time (Figure 3B).

Using DAF-2Da as a fluorescent probe for NO detection, we observed that treatment of RAW264.7 monocytes with LPS (500□ng/mL; used as a positive control of iNOS activation) for 24 h led to a significant increase in fluorescence, indicating elevated NO levels. Co-treatment with NPs composed of ProCIP and CMC at 30□µg/mL significantly reduced this fluorescence compared to LPS treatment alone (p<0.001; Figure 3C). NPs (made from ProCIP and CMC) were selected for cellular assays to avoid potential cytotoxicity and interference with iNOS expression caused by Cy5.5.

## 4. Conclusion

In this study, a protease-responsive polymeric iNOS-inhibiting prodrug, ProCIP, was successfully synthesized and formulated into nanoscale polyionic complexes through electrostatic assembly with anionic polymers. The resulting nanoparticles exhibited favorable physicochemical properties, and sustained release behavior, while demonstrating accelerated payload release in the presence of proteolytic enzymes. Importantly, the prodrug remained inactive under normal conditions but underwent enzyme-mediated activation, leading to substantial inhibition of iNOS activity. Cellular studies further confirmed efficient NP uptake and a significant reduction in intracellular NO production in LPS-stimulated macrophages. Collectively, these findings reveal the potential of ProCIP NPs as a protease-responsive platform for localized iNOS inhibition, offering a promising strategy to enhance therapeutic efficacy while minimizing systemic adverse effects associated with conventional iNOS inhibitors.

## Acknowledgements

This project receives funding from the European Union’s Horizon 2020 research and innovation programme under the Marie Skłodowska-Curie Grant Agreement No 101034324, as well as a grant from the Jaumotte-Demoulin Foundation. Some of the equipment used in this study was financed, in whole or in part, by the Walloon Region through the Technology Platforms of Excellence: ‘Alternative to animal experimentation’ and ‘Biogreen’. MA benefited from Grant n° 72290 from Tabriz University of Medical Sciences to support her Ph.D. thesis.

## Conflict of Interest

The authors declare no conflict of interest.

## Notes

### Competing Interest Statement

The authors have declared no competing interest.

